# Telocytes regulate macrophages in periodontal disease

**DOI:** 10.1101/2021.06.03.446871

**Authors:** Jing Zhao, Paul T. Sharpe

## Abstract

**Background:** Telocytes (TCs) or interstitial cells are characterised in vivo by their long projections that contact other cell types. Although telocytes can be found in many different tissues in luding the heart^1^, lung^2^ and intestine^3^, their tissue-specific roles are poorly understood. Here we identify a cell signalling role for telocytes in the periodontium whereby telocytes regulate macrophage activity.

**Methods:** We performed scRNA-seq and lineage tracing to identify TCs in mouse periodontium in homeostasis and periodontitis and carried out HGF signalling inhibition experiments using Tivantinib.

**Results:** We demonstrated that TCs are quiescent in homeostasis, however, they proliferate and serve as a major source of HGF in periodontitis. Macrophages receive telocyte-derived HGF signals and shift from an M1 to a M1/M2 hybrid state.

**Conclusions:** Our results reveal the source of HGF signals in periodontal tissue and provide new insights into the function of TCs in regulating macrophage behaviour in periodontitis through HGF/c-Met cell signalling, that may provide a novel approach in periodontitis treatment.

**Graphic abstract:** A population of telocytes (TC) have been identified in the periodontium. They are quiescent in homeostasis, however, in periodontitis, they are activated and send HGF signals to LPS activated iNOS+ (M1) macrophages. Macrophages receive the HGF signals via c-Met, which results in the expression of M2 marker Arg1, representing in an increase of M1/M2 hybrid macrophages. The expression of Arg1 can be inhibited by a HGF/c-Met selective inhibitor Tivantinib.

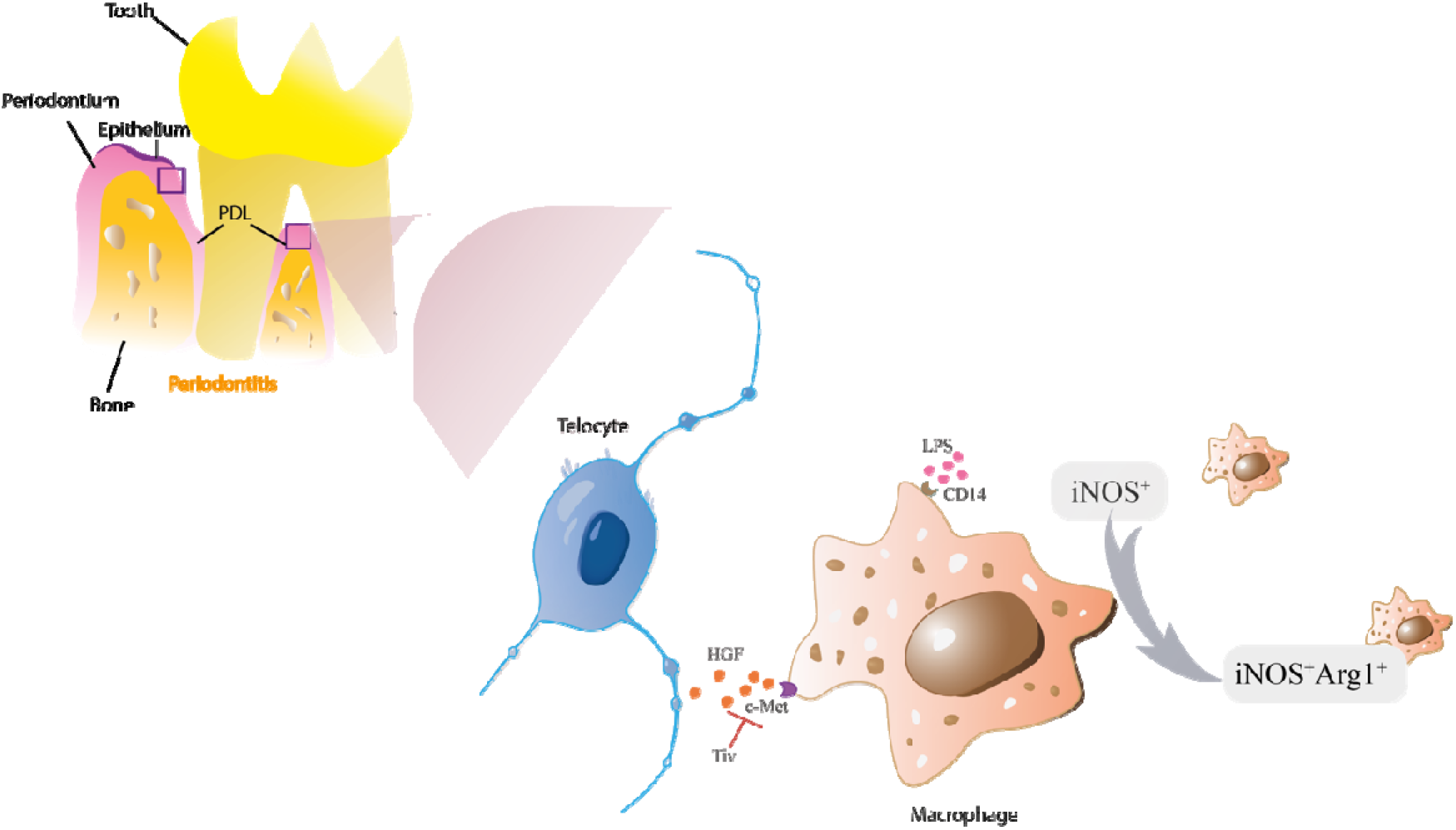

## Introduction

Periodontitis is an inflammatory disease of the periodontal ligament, the tissue that connects teeth to alveolar bone. It is a prevalent, incurable and continuous degenerative disease that results in bone loss and tooth loss. Individuals with periodontitis often exhibit gingiva recession, bleeding and tooth mobility^4^. All periodontitis (with attachment loss) develops out of gingivitis (no attachment loss) with poor prognosis but not all gingivitis develops into to periodontitis.

Numerous studies have focussed on the causes of periodontitis. Briefly, pathogens accumulate on the tooth surface (forming plaque), invade the periodontium tissue and release LPS (lipopolysaccharide), that results in inflammation and immunologic events. LPS causes the polarization of pro-inflammatory macrophages (M1). M1 macrophages release cytokines such as TNF-α,IFN-γ, IL-6 and IL-12, that contribute to the development and progression of inflammation-induced tissue destruction^5–8^. Hence, understanding the regulation of the inflammatory response is critical to understanding and treating periodontitis. The periodontal ligament (PDL), albeit only present as a thin layer, contains several different cell types. However, the understanding of cell populations in the PDL, their interactions, signalling pathways and how these are impacted upon by disease and inflammation are poorly understood.

In this study, we describe a cell type not previously identified in the PDL namely telocytes (interstitial cells). Telocytes are an enigmatic interstitial cell type that are best characterised by their unusual morphology having very long projections that make direct contacts with other cells. They are believed to a play a role in direct cell-cell communication establishing three-dimensional networks guiding tissue organization, mechanical sensing, regulating immune responses and phagocytic-like properties ^9,10^. Transmission electron microscopy (TEM) of TCs show extracellular vesicles bulging out from their membranes suggesting active physical communication with other cells^11^. To date possible cell signalling roles played by telocytes have not been fully described. It is only known that in the intestine, sub-epithelial telocytes are identified as an important source of Wnt signals to maintain the proliferation of intestinal stem cells ^3^.

TCs are identified in tissues by dual immunolabelling, most commonly CD34+/CD31−, CD34+/c-Kit+, CD34+/Vim+, CD34+/PDGFRα+ ^10,12–15^. In this study, we use CD34+/CD31− to identify PDL-resident TCs combined with genetic lineage tracing and sc-RNA sequencing to determine the role of TCs in homeostasis and periodontitis. We show that quiescent telocytes located near blood vessels are activated in periodontitis and regulate macrophages via the HGF/c-Met signalling pathway. The transition of macrophages provides possible therapeutic strategy for treating periodontitis.

## Methods and Materials

### Mice

All mice were maintained in the Biological Service Unit, New Hunts House, King’s College London. Mice were exposed to a 12◻hour light◻dark cycle and with food and water available ad libitum. Wild type CD1 mice were obtained from CRL (Charles River Laboratory, UK), *Wnt1^Cre/+^;R26^mTmG/+^* mice were from JAX 003829 and 007576 respectively. CD34^creERT2/+^; R26^tdTomato/+^ was a kind gift from Prof. Qingbo Xu (King’s College London). Three intraperitoneal injections of tamoxifen were given at a dose of 2 mg/30◻gbw (Sigma, T5648) for 3 consecutive days. Mice were sacrificed by exposure to a rising concentration of carbon dioxide or cervical dislocation followed by tissue dissection and tissue processing. All mouse work was approved by UK Home Office under the project license 70/7866 and P5F0A1579, approved by the KCL animal ethics committee.

### Animal disease model

Animals older than 8◻weeks were used to induce periodontitis. Mice were anaesthetized with Ketavat and Domitor, injected 10 mL/kg◻ i.p. The ligature procedure was performed as described^16^. Briefly, 5◻0 wax coated braided silk suture (COVIDIEN, S◻182) was tied around the upper second molar in order to induce periodontitis. Samples were collected at desired time points.

### HGF/c-MET pathway inhibition

CD1 mice were used to induce periodontitis. Tivantinib (130mg/kg in corn oil with 2.5% DMSO) was orally applied at day 5 post procedure. A control group was given corn oil with 2.5% DMSO. Samples were collected 12 hours later (n=3).

### Immunofluorescence

Maxillae were dissected and fixed in 4% PFA overnight. Samples were decalcified in 19% EDTA until soft enough to cut (~7days). Processed samples were then dehydrated with sucrose followed by embedding in OCT. Cryosections were fixed by 4% PFA. Sections were then subject to permeabilization by 0.2% Triton X◻100 (Sigma, X100), heat◻induced antigen retrieval, and blocking with 3% BSA. Sections were stained by the following antibodies: anti◻RFP (Abcam, Ab62341), anti -CD34 (Abcam, Ab81289 and Ab8158), anti ◻CD31 (Abcam, Ab7388 and Ab24590), anti ◻GFP (Abcam, Ab13970), anti -Arg1 (Abcam, Ab92274), anti -Met (Abcam, Ab51067), anti -HGF (Abcam, Ab83760) and anti-Ki67 (Abcam, Ab16667). Secondary antibodies included Alexa Fluor 488 (Invitrogen, A11039), Alexa Fluor 568 (Invitrogen, A11077), Alexa Fluor 633 (Invitrogen, A21052) and Alexa Fluor 488 (Invitrogen, A11008). Tyramide signal amplification (NEL744001KT, PerkinElmer) was performed for weak signals. Hoechst 33342 (Invitrogen 62249, 1:500) was used for DNA staining. Slides were mounted using Citifluor AF1 (EMS, 171024◻AF1) and cover◻slipped for microscopy. Zeiss Apotome or Leica TCS SP5 systems was used for acquiring images. Image J and Adobe Photoshop were used for image processing.

### Single-cell RNA sequencing and analysis

For scRNA-seq, adult CD1 mice were used. CD1 mice were sacrificed and dissected under a stereomicroscope with the gingiva carefully removed. Teeth were extracted and only the intact molars were kept. For periodontitis, only the second molars were used for subsequent use. The harvested molars were pooled and dissociated with 3 U/mL Collagenase P (COLLA◻RO, ROCHE) followed by incubation for 45◻minutes in a 37°C shaking water bath. The dissociation process was aided by dispersion with a 1 mL pipette every 15 minutes. Cells were then passed through a 40◻μm strainer (Falcon 352340) followed by FACS sorting for alive cells. Single cells in PBS with 0.04% ultrapure BSA were processed following a standard 10x genomic protocol (Chromium Single Cell 3’ v3). Count matrices were generated from the fastq files via CellRanger pipeline using Ensembl 97 genome annotation. Ambient and background RNA from the count matrices were first removed using CellBender remove◻background tool^17^. Cells express >1,000 features and with less than 20% mitochondria gene content were kept. 2270 cells were used for analysis. Batch effect was removed by the Seurat CCA approach^18^. Integrated data were subsequently scaled and PCA was performed. 30 dimensions were calculated based on variable features followed by UMAP^19^ for embedding and Louvain^20^ clustering (resolution 1) on knn graph. Macrophages (613 cells) were selected and re◻clustered. RNA velocity data were generated using the velocyto tool^21^. For gene enrichment analysis, metascape^22^ was used: genes highly expressed in TC cluster with avg_logFC >0 were selected, 863 input gene was used. Finally, cell-cell communication was estimated by using CellChat^23^. The communication between cell types was analysed based on the secreted signalling database.

### In vitro studies

PDL cells from CD1 mice (n=3) or *Wnt1^Cre/+^;R26^mTmG/+^* mice (n=3) were collected for cell culture. Tissues were treated as above to harvest single cell suspension. DMEM/F12media (3:1) supplemented with 20% FBS, L- glutamine and P/S was used for cell culture. Cells at passage 1 were used for analysis.

### Microcomputed tomography

Maxilla samples were fixed in 4% PFA overnight followed by three washes in PBS. Samples were scanned on a SCANCO μCT50 scanner with 70kVp voltage and a tube current of 114◻μA at 6μm isotropic voxel size. Scans were analysed by MicroView software.

### Statistical analysis

Statistical analysis was performed using an unpaired Student’s t-test using GraphPad Prism software. P < 0.05 were considered statistically significant.

## Results

### ScRNA-seq analysis reveals a telocyte population in PDL

The PDL is made up of both neural crest-derived and mesodermal-derived cell types and we have previously described the constituent cell populations using single cell transcriptomics^14^. This analysis compared adult PDL in homeostasis with PDL tissue from a ligature-induced periodontitis mouse model. The two datasets were integrated by performing a canonical correlation analysis (CCA)^24^ identifying 2,270 cells for analysis after filtering. These cells were further divided to 18 unsupervised clusters for annotation (Fig1-a). A small population of cells were found in the mesenchymal cell population that expresses CD34, however, they did not express the endothelial cell marker CD31 and we thus identified these cells as telocytes (TCs) (Fig1-b, SuppFig1).

**Fig 1.**
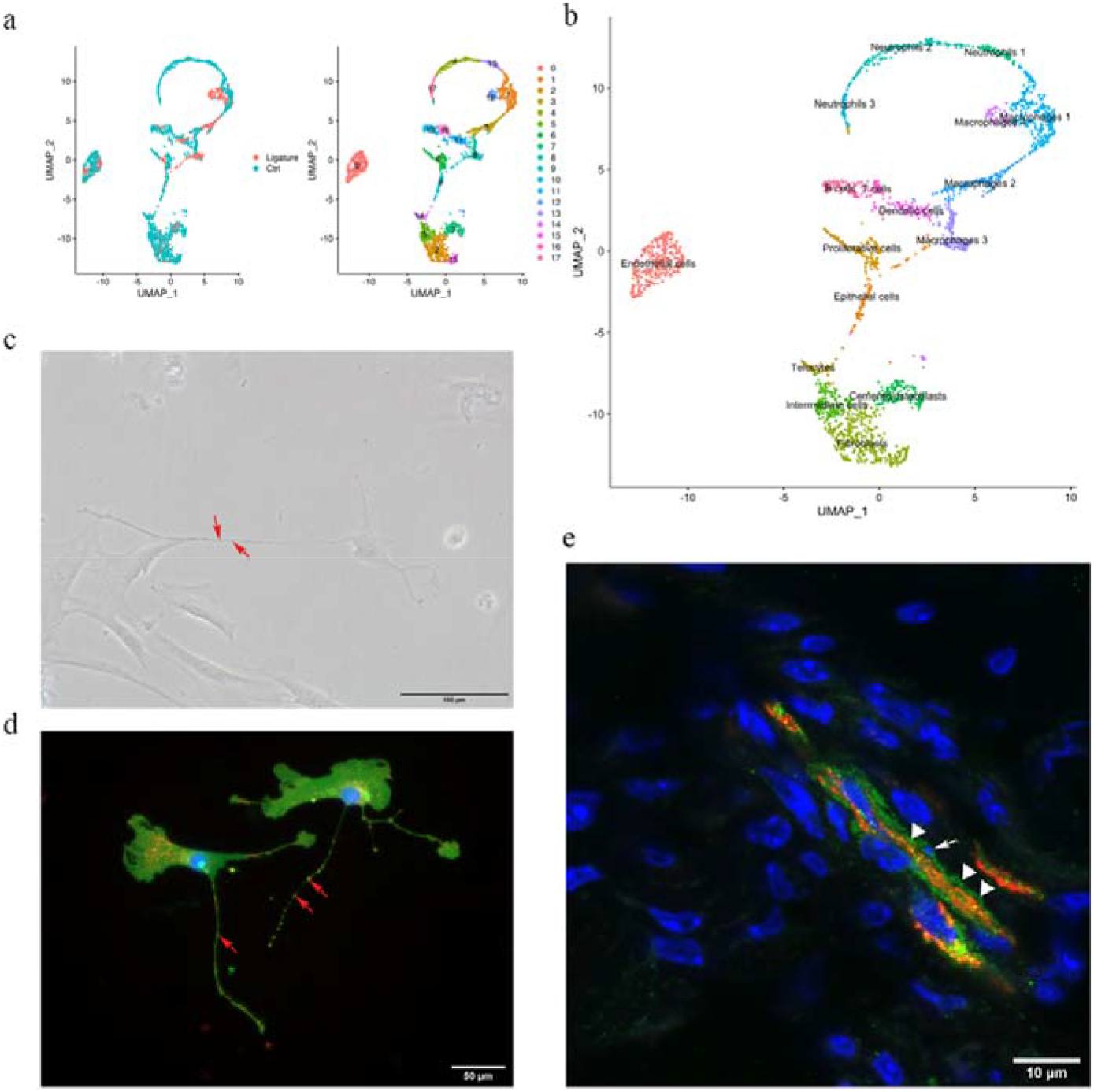
Telocytes in the PDL. **a.** PDL single cells from control mice and ligature treated mice were combined and clustered into 18 clusters. **b.** Identification of each cluster. Telocyte cluster were identified by CD34+ and CD31−. Macrophages are in four clusters. **c.** In vitro cell culture with CD1 PDL cells at passage 1 show characteristic telocyte structure, including podoms (red arrows) and podomers (between two arrows). **d.** Wnt1 lineage traced cells (GFP in green) were cultured and stained with CD34 in red, they show piriform cell body and moniliform podoms (red arrows) and podomers. **e.** Telocytes (CD34+CD31−) was detected near blood vessel (CD34+CD31+) in vivo, CD34 in green, CD31 in red. White arrow indicate the small nuclei and white triangle show the elongation of telocyte respectively.

To confirm these cells as TCs we first cultured PDL cells in vitro and searched for cells with a typical telocyte morphology. Typical TC cell morphology was observed with small cell bodies and a long cellular process called telopodes (Fig1-c). Telopodes consist of dilated portions (podoms) and thin segments in between (podomers). To determine if the TC-like cells are derived from neural crest, we collected PDL cells from *Wnt1^Cre/+^;R26^mTmG/+^* mice and stained for GFP and CD34. GFP positive cells showed TC structures with podoms and podomers, which were also positive for CD34 (Fig1-d). To identify TC location *in situ*, we co-immunostained sections with CD34 and CD31. As shown in Fig1-e, TC cells (CD34+/CD31−) were found in close association with blood vessels (CD34+/CD31+), had small cell nuclei and long cell protrusions. These data collectively suggest that CD34+/CD31− telocytes are present in PDL located in the vicinity of blood vessels.

### Quiescent telocytes are activated in periodontitis

Since telocytes only account for a small subset of cells in PDL, we asked whether these cells are quiescent or actively proliferating during homeostasis. Lineage tracing using CD34^creERT2/+^; R26^tdTomato/+^ mice followed by a 1d-1yr post-tamoxifen chase period revealed that CD34+/CD31− cell numbers did not increase to any significant extent (Fig2), suggesting TCs are a small, quiescent cell population in homeostasis. In order to investigate if TCs responded to disease we used our established ligature-induced periodontitis mouse model where sutures are placed around the second molars^14^. The ligature leads to plaque accumulation and thus facilitates the invasion of bacteria^25^. By measuring the distance between alveolar bone crest and cemento-enamel junction (ABC-CEJ distance) (Fig3-a), we found that bone loss reached a maximum between day 4-7 for all three molars (Fig3-b). Notably, even though only the second molar was subjected to a ligature, the first and third molars also showed some bone loss at early time points, suggesting they are affected by the ligature-induced periodontitis to some extent. However, the first and third molar bone loss was recovered at longer time points (Fig3-b), indicating the self-recovery ability from milder periodontitis.

**Fig 2.**
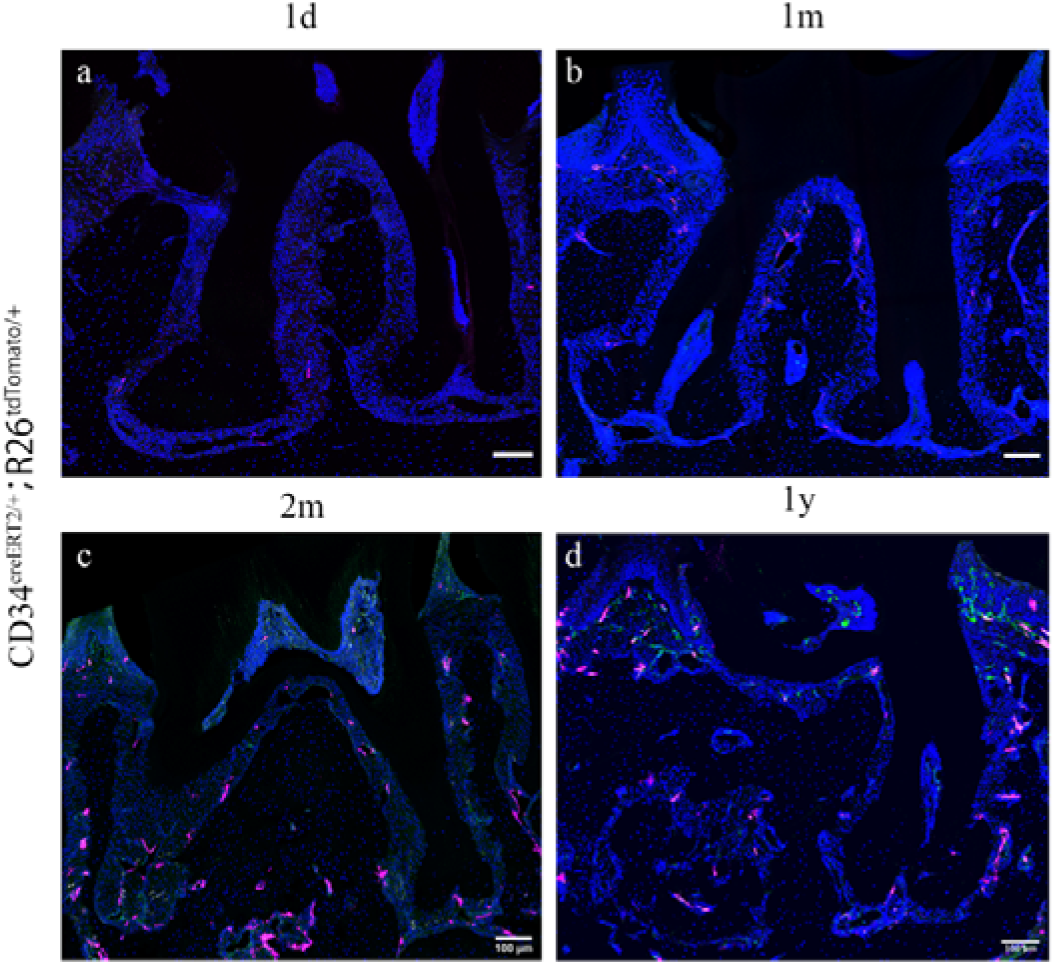
CD34^creERT2^ lineage tracing showing limited contribution of telocytes to PDL homeostasis in adulthood. In the adult stage, CD34^creERT2/+^; R26^tdTomato/+^ mice were used to trace from 7 weeks for 1 day (**a**), 1 month (**b**), 2 months (**c**) and 1 year (**d**). CD34 lineage traced cells in red, CD31 were co-stained in green. Increase of cell number was not detected as the extending of tracing time. Scale bars = 100 μm.

**Fig 3.**
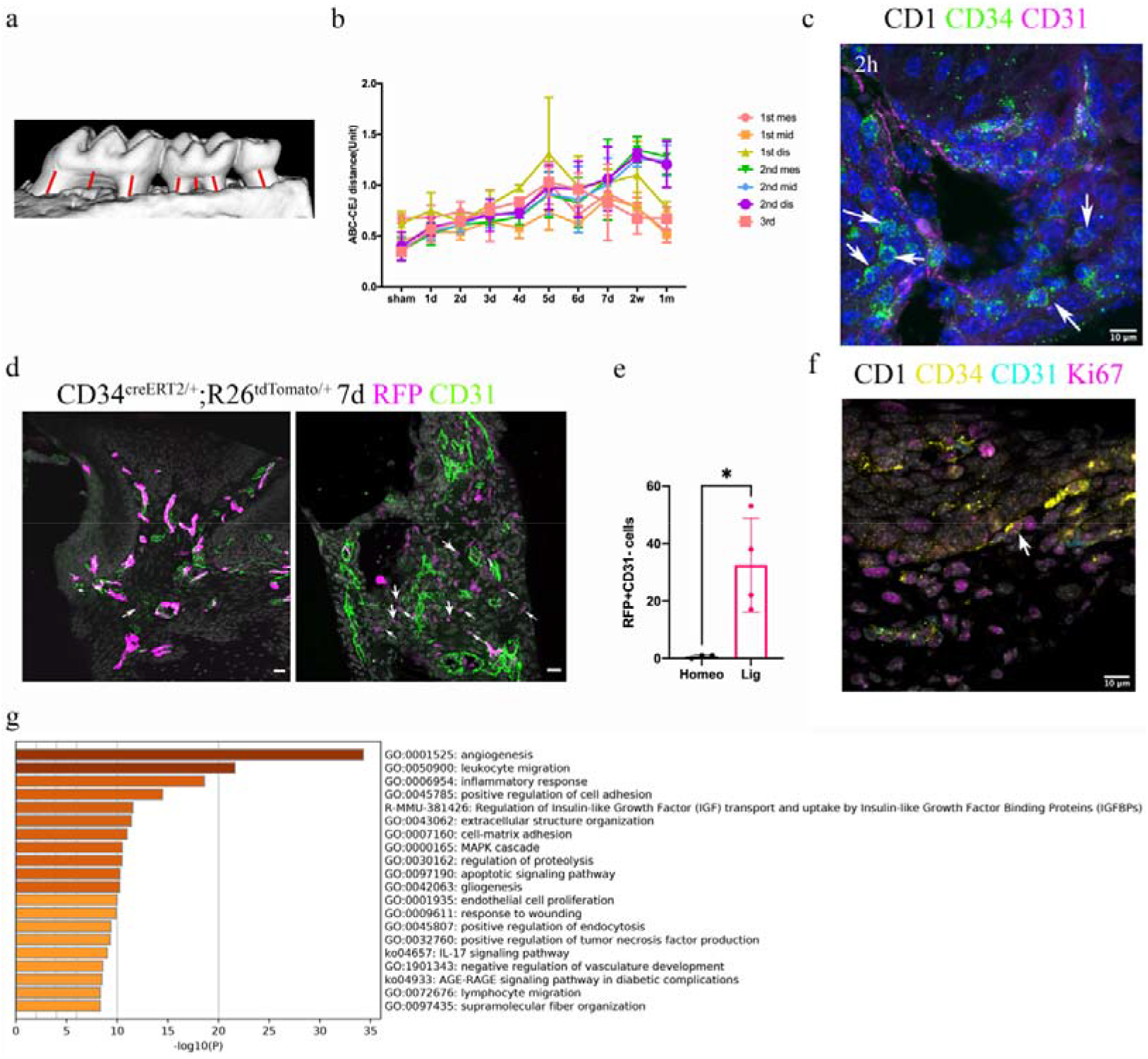
Telocytes are activated in response to periodontitis by increasing their number. **a.** Illustration of bone loss measurement. For the first and second molars, the ABC-CEJ distance of mesial and distal roots in parallel to the root long axis and the ABC-CEJ distance in the trifurcation area were measured, for the third molar, the ABC-CEJ distance in the middle of the tooth was measured. **b.** Quantification from micro CT results indicates the change of bone loss in periodontitis plotted by time course (n=3). Hard tissue around the second molar is severely affected, the time for all molars reaching the bone loss plateau is between day5 to day7. 1 unit = 0.4mm (n=3). **c.** An accumulation of telocytes was found in periodontitis as early as 2 hours after the ligation procedure. These telocytes (CD34, green) were mostly found around blood vessels (CD31, red). Scale bars = 10μm. **d.** CD34^creERT2/+^; R26^tdTomato/+^ mice were given three tamoxifen injections started from the procedure day and harvested at day 7. Significantly increased CD34+ cells (red) were observed in the periodontitis group. There were more endothelial cells (green) observed at day 7 of periodontitis. The telocyte (red) derived cells were not overlapping with endothelial cells (green). Scale bars = 20μm. **e.** Statistical analysis of numbers of telocytes comparing homeostasis (n=3) and periodontitis (n=4). **f**. Telocytes (CD34+, CD31−) express a proliferation marker, Ki67 in periodontitis. CD34: yellow, CD31: cyan, Ki67: magenta, nuclei: grey. **g.** 863 input genes highly expressed in TC cluster with avg_logFC >0 were selected for gene enrichment analysis. The 20-best p-value terms was plotted. The bar plot is colored by p-values.

We visualised TCs with CD34 and CD31 antibodies in ligature PDL at different time points. After 2h, there was an obvious increase in the number of CD34+/CD31− TCs close to blood vessels (Fig3-c, Supp.Fig2). CD34^creERT2/+^; R26^tdTomato/+^ mice tracing for 7 days also showed an increase in tdTomato+/CD31− cells in periodontitis (Fig3-d,e). These TCs also expressed the proliferation marker Ki67 (Fig3-f). These data indicate that TCs proliferate in the PDL following ligature-induced periodontitis.

It has been demonstrated that TCs can secrete extracellular vesicles^11^, suggesting they may have a role in cell signalling. Gene enrichment analysis of the single cell RNA-seq datasets identified angiogenesis, leukocyte migration and inflammatory responses as the three top pathways in PDL TCs (Fig3-g).

### Telocytes regulate macrophages via HGF/c-Met signalling pathway

To functionally understand the differences between homeostasis and periodontitis, we interrogated the RNA-seq datasets to compare the two conditions with respect to cell-cell communication pathways. We identified the CHEMERIN, HGF, IFN-I, IL16, LIFR and APELIN pathways as not being active during homeostasis but active in periodontitis (Fig4-a). Telocytes were found to express *Hgf* and *Flt3* (Fig4-b), highlighting a potential role for the HGF signalling pathway.

**Fig 4.**
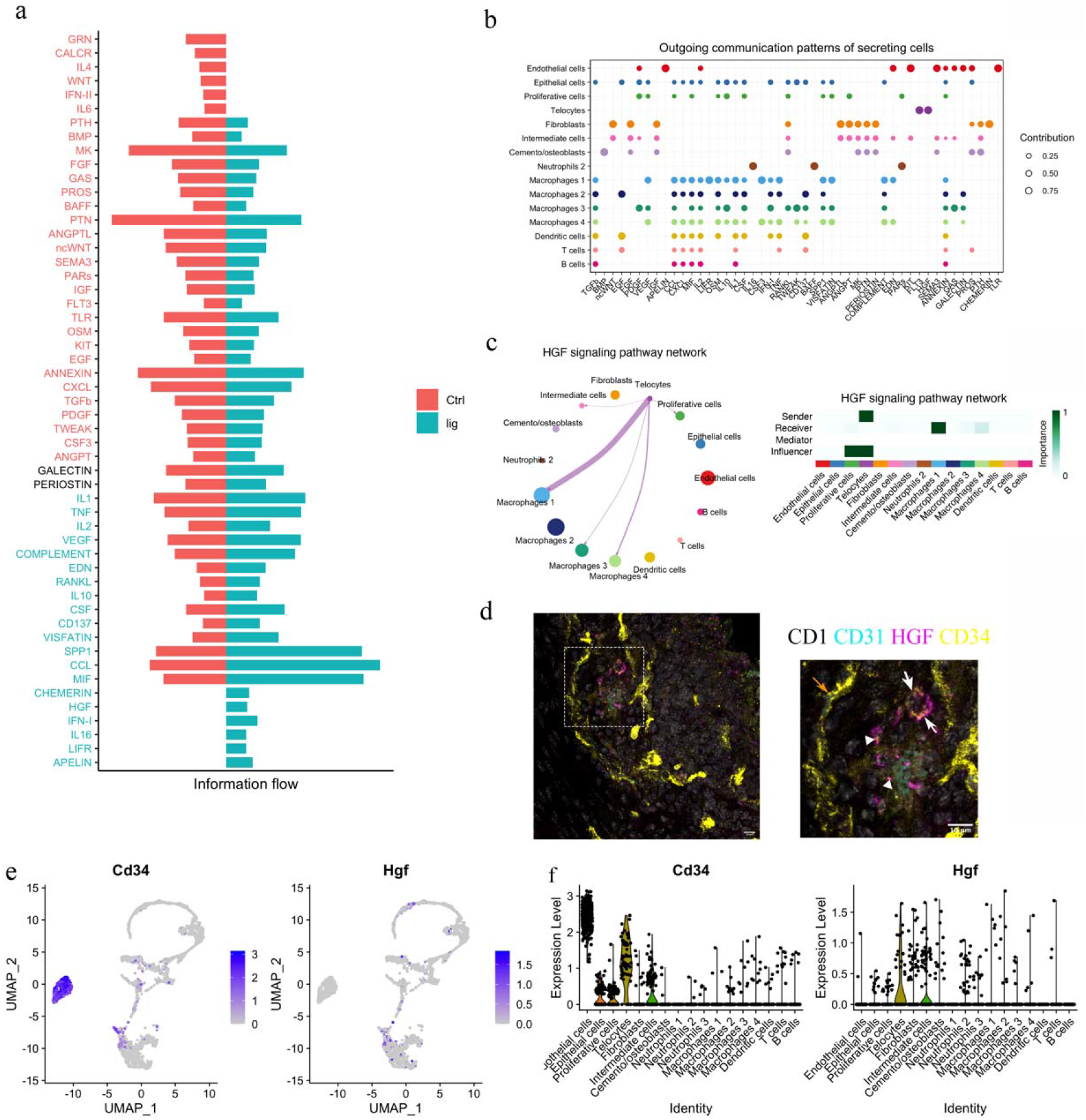
Telocytes regulate macrophages via HGF/c-Met signalling pathway. **a.** Comparison analysis show HGF pathway is upregulated in periodontitis **b.** Cell-cell communication analysis was performed on disease dataset based on secreted signals database. The outcoming patterns were plotted. Bubble plot suggest that telocytes send signal molecules in FLT3 and HGF signalling pathways exclusively in periodontitis (dark purple dots). **c.** Circle plot and heatmap suggest telocytes send HGF signals to macrophage clusters 1, 3, 4 in periodontitis. **d.** Immunostaining on CD1 mice periodontitis tissue for CD34+/CD31− cells indicate telocytes (white arrows) and CD34+/CD31+ cells for endothelial cells (orange arrow) from CD1 mice. The typical morphology of telocytes, podoms are denoted with arrow heads. The telocytes were expressing HGF (magenta). Scale bars = 10μm. **e.** Expression of Cd34 and Hgf were plotted. **f.** Cd34 is expressed in endothelial cells and telocytes, Hgf is expressed in telocytes but not endothelial cells. and some intermediate cells that close to telocytes.

HGF (hepatocyte growth factor) was originally found in liver as a potential hepatocyte mitogen^26^. It is involved in repair and regeneration as a healing factor^27–29^. In mouse ligature-induced periodontitis, our RNA-seq cell-cell communication analysis identified macrophages as a target of HGF signalling from TCs (Fig4-c). Of the 4 macrophage clusters identified, TCs are identified as potentially interacting with macrophage clusters 1 and 4. To confirm that TCs express HGF, triple immunofluorescence staining was performed with CD34, CD31 and HGF. Endothelial cells (CD34+/CD31+) did not express HGF, whereas TCs (CD34+/CD31−) with telopodes expressed HGF (Fig4-d). This is thus consistent with sc-RNA seq analysis (Fig4-e,f). In conclusion, telocytes express HGF and are the only cells that send this signal in periodontitis. The recipient cells are macrophages which express the HGF-receptor HGFR (Hepatocyte growth factor receptor, encoded by *c-Met*).

### Macrophages receiving HGF signals show M1-M2 transition

Four clusters of macrophages were identified from the single cell transcriptomics analysis (Fig. 5). DEG (differentially expressed genes) analysis suggested macrophage cluster 1 express *Acod1*, *Trem1*; macrophage cluster 2 express *Mc4a4c*, *Ccr2*; macrophage cluster 3 express *Aif1*, *Mrc1*; macrophage cluster 4 express *Cd36*, *Arg1* (Fig5-a). In order to gain a better understanding of these cells, we performed unsupervised clustering on the 4 clusters. Macrophages from homeostasis (right) and periodontitis (left) are clearly separated (Fig5-b).

**Fig 5.**
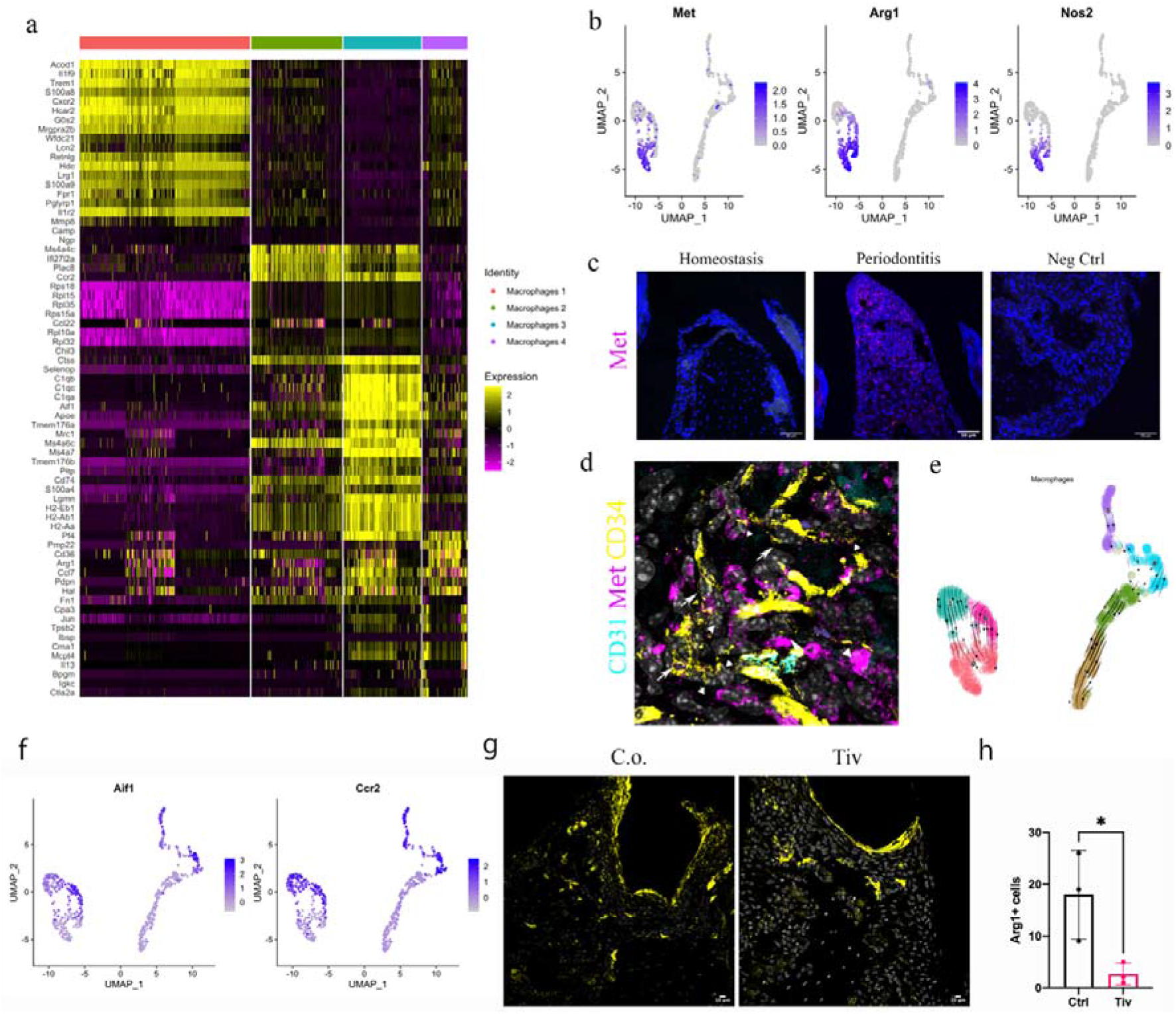
HGF/c-Met signalling drives M1 to M2 transition. **a.** Heatmap presenting the DEG of 4 macrophage subpopulations. **b.** Macrophages was extracted from the complete dataset and re-clustered. All the macrophages in disease dataset are in the left cluster. Feature plots show c-Met expressing macrophages express M1 marker *Nos2* and M2 marker *Arg1*. **c.** Met protein was not detected in PDL homeostasis but in periodontitis. Met: magenta, nuclei: blue. Scale bars = 50μm. **d.** Telocytes (yellow, indicated by arrows) are making contact with Met expressing cells (magenta) by using their protrusions (telopodes indicated by arrow heads). CD34: yellow, CD31: cyan, Met: magenta, nuclei: grey. Scale bar = 10μm. **e.** RNA velocity show, *c-Met*+ macrophages derive from cluster 3 and 6. **f.** Macrophages expressing *c-Met* may derive from cluster 6 (*Ccr2*+*Aif1*^hi^) and cluster 3 (*Ccr2*+ *Aif1*^lo^) macrophages. **g.** CD1 mice were used to induce periodontitis. Corn oil or Tiv were given once at day 5 post procedure. Samples were collected 12 hours after drug delivery. Mice given corn oil (left) show more cells express ARG1+ cells (yellow) in periodontium than the mice given Tivantinib (right). Scale bars = 10μm. **h.** Statistics analysis suggest significance difference of ARG1 expression between ctrl group and Tivantinib treated group (p<0.05).

The HGF signalling pathway has a sole ligand-receptor pair, HGF and HGFR (c-Met). Interestingly, macrophages expressing c-*Met* also expressed *Arg1* (encodes Arginase1, ARG1) and *Nos2* (encodes iNOS, inducible nitric oxide synthase) (Fig5-b).

In periodontitis, iNOS mediates the pathological effect of LPS and it is a marker of inflammatory M1 macrophages, which is an indicator of disease progression. *Arg1*, a gene expressed in M2 macrophages, however, is believed to decrease LPS-induced pro-inflammatory cytokine production^30^. Therefore, macrophages activated by HGF signals in the PDL can be considered to be in a hybrid state. Previous studies indicate that monocytes in an inflammatory environment first polarize to M1, then M2 subject to micro-environmental signals^31,32^. It is well known that ARG1 competes with iNOS for their common substrate L-arginine^30,33^. Moreover HGF/c-Met is reported to induce *c-Met*+/*Nos2*+ macrophages to an M2-like phenotype via overexpression of *Arg1* ^34^. These data collectively suggest that the increase in TCs numbers mediates a transition from the M1 state to the M2 state in periodontitis.

During homeostasis, macrophages in PDL do not express c-Met protein, however, macrophages start to express c-Met in periodontitis (Fig5-c). C-Met expressing cells are located in close proximity to the telopodes of TCs (Fig5-d), indicating TCs likely make physical contact with c-Met+ macrophages. Thus, it is conceivable that TCs promote the overexpression of *Arg1* in periodontitis which further leads to the hybrid state of macrophages. Additionally, RNA velocity analysis suggests that these *Arg1*+/*Nos2*+ macrophages derive from *Ccr2*^hi^*Aif1*^lo^ (cluster 3) cells and *Ccr2*^lo^*Aif1*^hi^ cells (cluster 6) (Fig5-e, f).

To experimentally determine if *Arg1* expression in macrophages is increased by HGF signals from TCs, tivantinib (ARQ197), a small molecule inhibitor was used to inhibit HGF/c-Met signalling. Tivantinib is a highly selective, non-ATP competitive, orally available inhibitor of c-Met ^35^. We observed that when tivantinib was administered to mice with ligature-induced periodontitis, Arg1+ cell numbers significantly decreased (Fig5-g, h), indicating that inhibition of the HGF-c-Met signalling interaction in macrophages blocks their polarisation.

## Discussion

Based on *in vitro*, *in vivo* and *in silico* studies, we identified a neural crest-derived cell population by morphology and expression of CD34+/CD31− in mouse periodontium. These cells, telocytes, are in a quiescent state in normal periodontal tissue unless challenged by periodontitis whereupon they increase in number and secrete HGF. Cleaved HGF can activate c-Met and downstream signalling pathways^36^. It is believed that pro-HGF, the inactive precursor is secreted and processed by HGF activator (HGFAC), a zymogen in the circulation to achieve its function ^37^.

We observed TCs, as a major source of HGF, making physical contact with macrophages. Macrophages receiving HGF signals express *c-Met*, *iNos* and *Arg1*, representing a M1/M2 hybrid state. The *Arg1*+/*Nos2*+ hybrid macrophages have been recently proposed to exist in other tissues ^38–40^. These cells do not exist in homeostasis but are detected in periodontitis identifying HGF/c-Met as a key macrophage regulatory pathway. Our findings are consistent with reports that LPS causes M1 polarization and a shift towards M2 polarization mediated by HGF signals^34^. Thus, expression of two competing enzymes, iNOS and ARG1 in c-Met expressing cells results from regulation by TCs, which shifts M1 macrophages to an M2 phenotype, resulting in a hybrid state. The transition can be effectively inhibited by an HGF/c-Met inhibitor, tivantinib. It is reported that the transition from LPS-induced M1 macrophages to M2 macrophages is controlled by PI3K or CaMKKβ-AMPK signalling pathway in c-Met expressing cells via induction of Arg1 expression^34,41^. We propose the underlying mechanism is possibly the activation of HGF/c-Met signalling pathway triggered the activation of PI3K or CaMKKβ-AMPK signalling pathway in macrophages.

The presence of M1 macrophages can cause bone loss whereas M2 macrophages can help prevent bone loss in the PDL. TCs showed the ability to shift M1 to M2, indicating that HGF secreted by TCs should be beneficial in reducing bone loss. By comparing the bone loss of the second molars with the first molars and third molars from 1 day to 1 month post ligature treatment, we found that only the molars without ligature treatment showed an ability to recover bone loss caused by periodontitis. Second molars, which have the ligature throughout, showed consistent bone loss indicating any ability of TCs to reduce bone loss is limited by constant physical stimuli, supporting the importance of maintaining oral hygiene in future clinical applications.

Ligature-induced periodontitis is considered as an appropriate model to mimic human periodontitis^42^. However, whether TCs are able to control the progression of human periodontitis requires further investigation. From the aspect of this study, an optimistic outcome is expected given the activation of TCs or HGF/c-Met pathway under careful maintenance of oral hygiene. Coincidentally, exogenous application with HGF was found to improve periodontal bone regeneration in swine^43^.

Additionally, TCs may have a role in angiogenesis as shown in the gene enrichment terms (Fig3-e). More CD31+ cells were noticed in periodontitis samples (Fig3-d). Vascularization is considered important for periodontal regeneration^44^. Therefore, the role of TCs in periodontitis may not only be the regulation of macrophages through HGF/c-Met signalling pathway but also through angiogenesis. TCs are the cells provide niche signals in intestine^3^. Future work may focus on the signals telocytes are sending to adjacent niche cells including stem cells^45^, endothelial cells^46^ and nerve cells^45^.

Collectively, our study demonstrates for the first time that TCs increase in number in periodontitis and communicate with immune cells to positively regulate periodontitis via HGF. The activation of HGF/c-Met signalling pathway or the exogenous use of activated TCs may be a promising therapeutic measure against periodontitis. This function of TCs may also present in other inflammatory disease where TCs exist such as arthritis^47^. Furthermore, our study also has implications for cancer research, where TCs were found present and HGF/c-Met signalling were found essential for cancer metastasis^48,49^.

## Data Availability

Sequencing data have been deposited in the GEO database under accession code GSE160358 and GSE167917.

## Author contributions

J.Z conceived, designed and performed the experiments and the analysis. P.S. supervised the research. J.Z and P.S. wrote the manuscript. P.S. approved the final version of the manuscript.

## Acknowledgements

We would like to thank Prof. Qingbo Xu (King’s College London) for kindly provide the CD34^CreERT2/+^; R26^tdTomato/+^ mice. We thank Dhivya Chandrasekaran, Fernanda Suzano, and Christopher Healy for technical assistance. We sincerely appreciate Dr.Cynthia Andoniadou for her valuable comments and suggestions, which helped us to improve the quality of the manuscript. The research described was supported by the National Institute for Health Research’s Biomedical Research Centre based at Guy’s and St Thomas’ NHS Foundation Trust and King’s College London. The views expressed are those of the authors and not necessarily those of the NHS, the National Institute for Health Research, or the Department of Health. Jing Zhao is supported by the China Scholarship Council.

---

**Supp. Fig-1.**
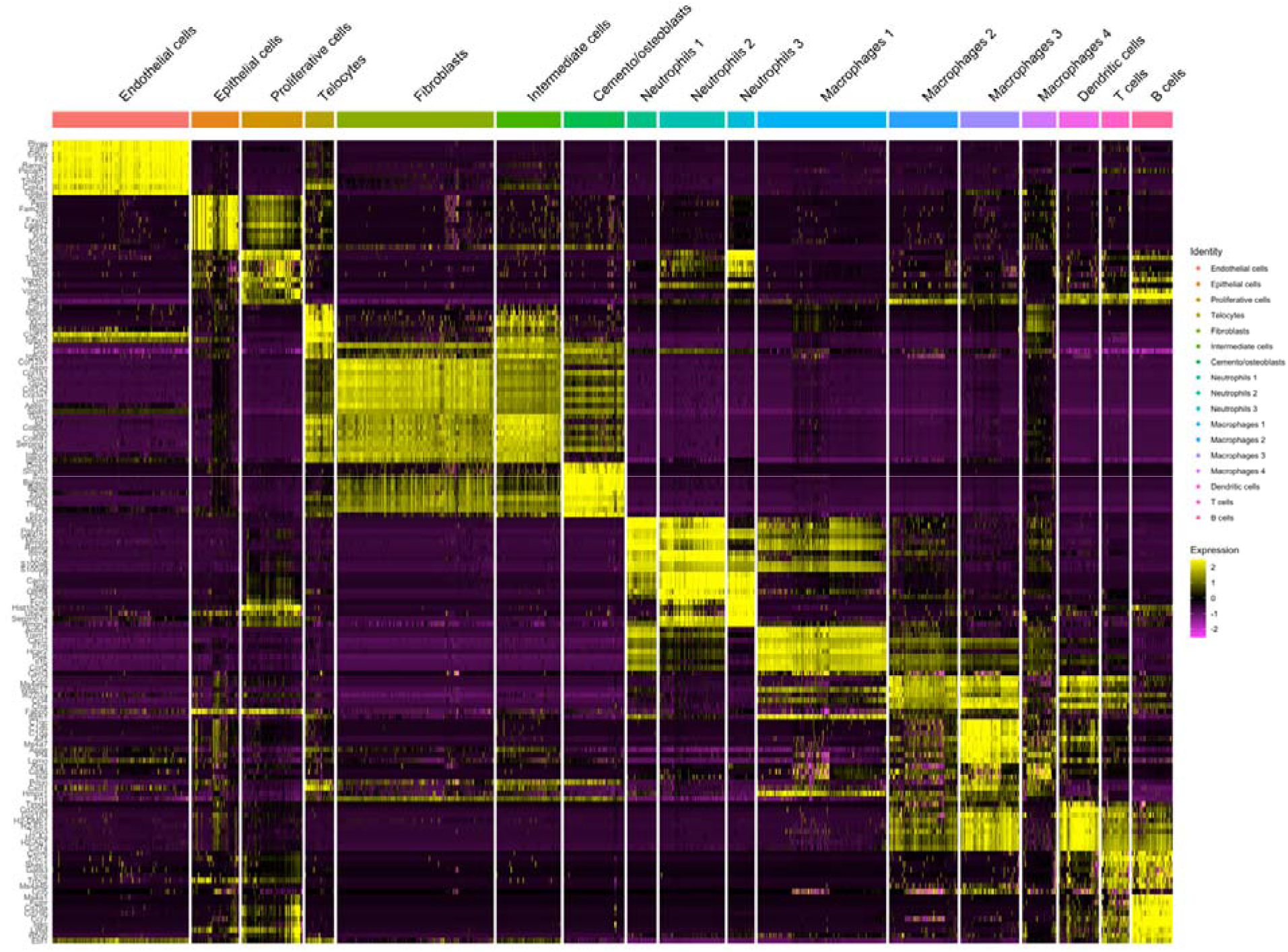
Heatmap of cell clusters show telocytes distinct from other mesenchymal cells.

**Supp. Fig-2.**
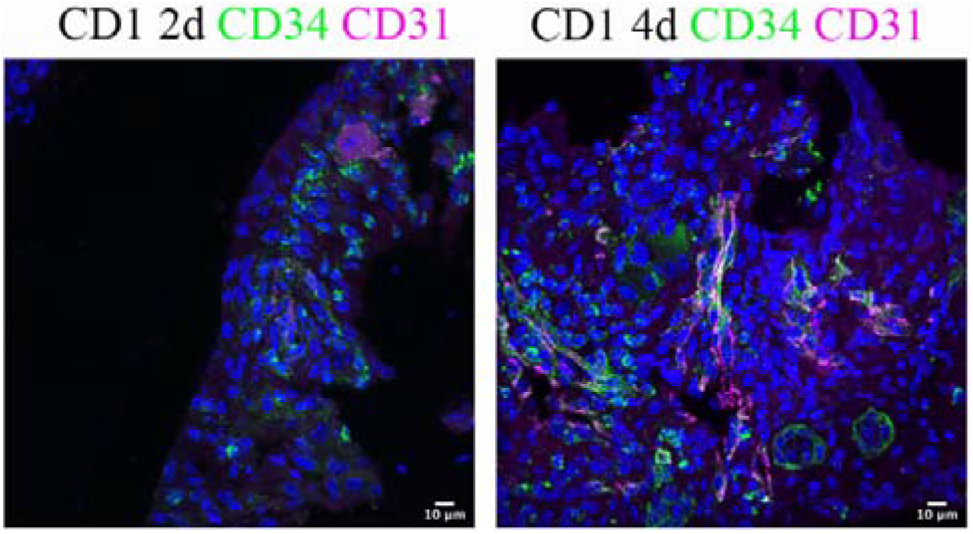
An increased number of telocytes was observed at day 2 and day 4 post procedure.

